# Resolving the molecular architecture of the photoreceptor active zone by MINFLUX nanoscopy

**DOI:** 10.1101/2021.05.28.446138

**Authors:** Chad P. Grabner, Isabelle Jansen, Jakob Neef, Tobias Weiss, Roman Schmidt, Dietmar Riedel, Christian A. Wurm, Tobias Moser

## Abstract

Cells assemble macromolecular complexes into scaffoldings that serve as substrates for catalytic processes. Years of molecular neurobiology indicate that neurotransmission depends on such optimization strategies, yet the molecular topography of the presynaptic Active Zone (AZ) where transmitter is released upon synaptic vesicle (SV) fusion remains to be visualized. Therefore, we implemented MINFLUX optical nanoscopy to resolve the AZ of rod photoreceptors. To facilitate MINFLUX nanoscopy of the AZ, we developed and verified an immobilization technique, we name Heat Assisted Rapid Dehydration (HARD). Here fresh retinal slices are directly stamped onto glass coverslips yielding a single layer of rod AZs. These AZs exhibited excellent labeling efficiency and minimal signal redundancy in the Z-direction. Our data indicate that the SV release site is a molecular complex of bassoon-Rab3-binding molecule 2 (RIM2)-ubMunc13-2-CAST. The complexes are serially duplicated longitudinally, and reflected in register along the axis of symmetry of the synaptic ribbon.

**One sentence summary:** Structural motifs formed by active zone proteins at the photoreceptor synapse.

---

The combination of multi-scale structural approaches with molecular neurobiology and electrophysiology has given us a dynamic view of the AZ. The classic electron microscopic (EM) experiment that motivated intense interest in AZ structure arose from images of freeze fractured frog neuromuscular junction^1^. In short, the study resolved two rows of paired intramembrane particles, running for microns in the presynaptic plasma membrane (PM), which were associated with stimulation-dependent synaptic vesicle (SV) fusion. However, the molecular identity of the intramembrane particles remained unknown. Inspection of this AZ in smaller volumes using high resolution EM tomography showed a pair of thin, electron dense tethers connecting the SVs to the PM (via a ‘rib’) and to the intra-membrane particles (via ‘pegs’)^2,3^. Efforts to resolve the physical positions of AZ proteins (molecular topography) have involved immuno-EM approaches to study several types of synapses^4,5^. These studies have put years of biochemical work into 2D maps of the AZ^6^ with a resolution between 10 to 20 nm. Unfortunately, the labeling density in immuno-EM is typically low^7^, rendering imaging of multiple proteins challenging – especially if more than one protein of interest should be imaged in a sample. Since several years, super-resolution light microscopy as stimulated emission depletion (STED) microscopy has offered a complementary view, with its greatest advantage being the ease of imaging multiple proteins at high resolution^8–12^. However, a disadvantage, compared to immuno-EM, is that spatial resolution has remained limited to ∼25 nm in the XY plane and the Z-direction is even less resolved. The outstanding goal has been to resolve the molecular topography of the AZ with a precision in the range of 1 to 10 nm using optical nanoscopy. This has recently become feasible by means of MINFLUX (minimal photon fluxes) nanoscopy that achieves a resolution of few nanometers with much lower need of photons than conventional localization microscopy approaches^13,14^.

As for electron tomography of presynaptic structure^2^, AZs featuring stereotyped patterns of repeated structural motifs are particularly suitable for the analysis of molecular AZ topography with optical nanoscopy^8,10–12,15,16^. Among these attractive targets are ribbon synapses which feature “ribbons”, i.e. large electron-dense projections emanating from the presynaptic membrane that tether SVs and are primarily made of the protein ribeye^17^. Rod photoreceptor presynaptic terminals (rod spherules) form very large ribbons with an average length of 1.5 µm in mice (Figure 1, review in ^18,19^). They express some of the same proteins localized to the presynaptic densities at conventional synapses, such as bassoon^20^, CAST^21^, rab interacting protein 2 (RIM2)^22,23^, and piccolino (a short splice variant of piccolo)^4,20,24^. Approximately 60 - 80 SVs are wedged between the ribbon’s base and the membrane^25,26^ and are likely release-ready, since a similar number of SV fuses within 1 ms upon strong depolarization^27^. Here we established a protocol for immobilizing rod spherules and used MINFLUX nanoscopy to reveal the molecular AZ topography.

**Figure 1.**
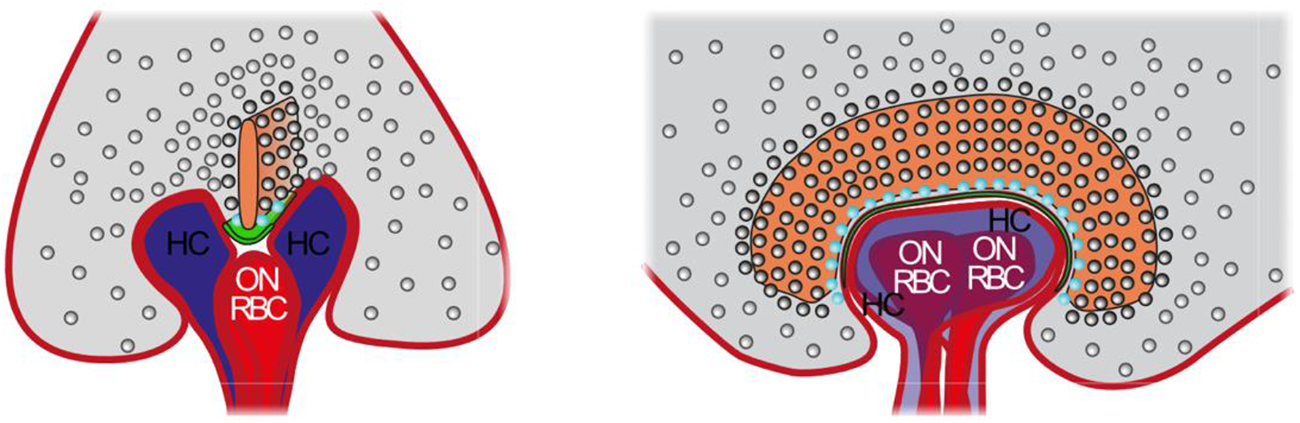
Morphology of the rod photoreceptor ribbon synapse. Schematic representations of the rod ribbon synapse in two perspectives: front and side views (left and right). The synaptic ribbon within the presynapse (rod spherule, gray) is drawn in orange, the presynaptic (arciform) density in green, and synaptic vesicles as gray spheres, except for membrane proximal vesicles (cyan). Modified from ref.18.

## Immobilizing a single layer of rod ribbons by Heat Assisted Rapid Dehydration

In order to make the rod AZ accessible to MINFLUX imaging and to reduce the background, we set out to establish a protocol for immobilizing rod spherules that are contained in the outer plexiform layer of the retina (Figure 2a, b) onto a planar substrate. For this we used fresh retinal slices, as prepared for electrophysiological recordings, and placed them directly onto heated glass cover slips for a few seconds, leading to the immobilization of a single layer of rods and rod spherules on the glass (Figure 2a). Heat Assisted Rapid Dehydration (HARD) i) avoids chemical fixation of the tissue, ii) does not require surface functionalization of the glass, and iii) makes the AZs readily accessible for immunolabeling without detergent permeabilization - probably via cracks in the membrane. Confocal imaging of HARD samples labeled for the ribbon specific protein ribeye readily identified rod ribbons (uniform 1.5 µm contour length vs. cone: 0.7 µm and bipolar 0.2 µm) that associated with other AZ markers (Figure 2b, c). As expected, the density of immunolabeled rod ribbons was much lower in the HARD sample than in the retinal slices (Figure 2b) typically providing for sufficient separation of the AZs for nanoscopic analysis. In addition, stacks of confocal sections confirmed the AZs were attached to the glass and contained within (Figure 2c, d) approximately 1 µm from the glass surface (Figure 2c, d). To investigate the transferred structures in terms of identity and structural integrity, we used electron microscopy. We readily identified photoreceptors containing characteristic sub-structures such as membrane discs of outer segments and kinocilia (Figure 2e). The structural preservation of the material is evidenced by the clear visibility of intact membrane structures and organelles as mitochondria (Figure 2e).

**Figure 2:**
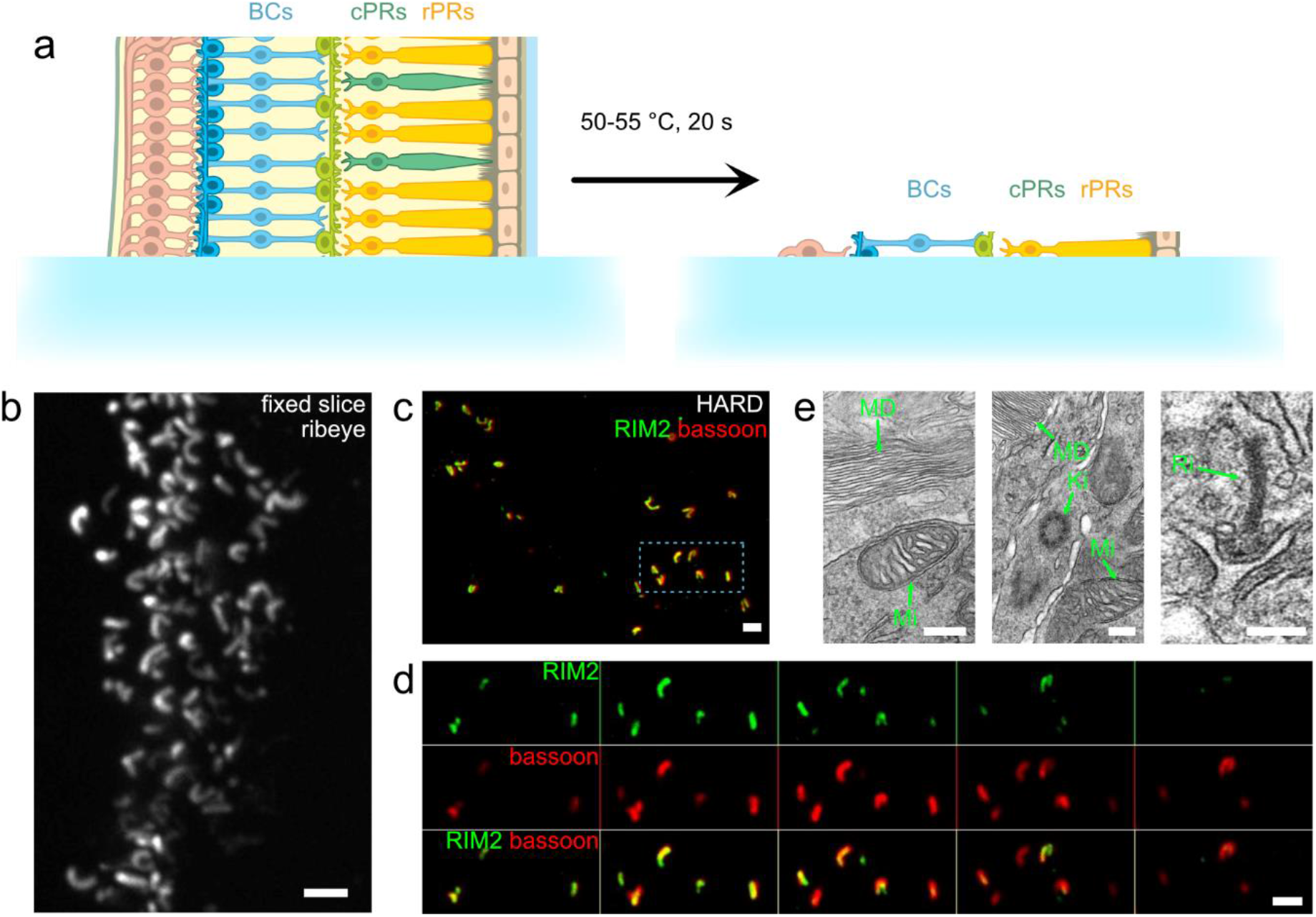
Immobilizing rod spherules onto coverslips by Heat Assisted Rapid Dehydration (HARD) (a) Schematic representations of the HARD procedure: acute 200 µm slices were placed on glass cover slips residing on a heat block (left) for transferring a cell layer onto the glass that remained after gently peeling off the slice after 20 seconds at 50-55 °C. rPRs: rod photoreceptors, cPRs: cone photoreceptors, BCs: bipolar cells, GCs: ganglion cells. Figure modified from ref. ^28^. (b) Exemplary projection of confocal sections of a PFA fixed retinal slice immunolabeled for the ribbon marker ribeye highlighting the rod photoreceptor ribbons in the outer plexiform layer of the retina. Scale: 2 µm. (c) Exemplary projection of confocal sections of a HARD sample immunolabeled for RIM2 and bassoon, markers of the presynaptic AZ. Scale: 2 µm. (d) confocal sections from the area boxed in (c) showing RIM2 (top) and bassoon (middle) in separation and merged (lower). Left to right: increasing height in z. Z-steps between images are 250 nm. Scale: 2 µm. (e) Exemplary transmission electron micrographs of a HARD sample showing membrane discs of outer segments (MD) and a mitochondrion (Mi; left panel), a kinocilium (Ki), membrane discs of outer segments (MD) and a mitochondrion (Mi; middle panel) and a photoreceptor ribbon synapse (Ri; right panel). Scale: 200 nm.

### Bassoon forms pairs of periodic spots along the longitudinal axis of the active zone

Bassoon is 420 kDa multidomain protein of the active zone, contributes to the presynaptic density and is critical for anchoring the ribbon to the presynaptic density^4,20,29–33^. Consistent with these findings, stimulated emission depletion (STED) nanoscopy showed bassoon lining the inner contour of the rod ribbon in HARD samples, when laying on their side (Figure 3a). We then moved to MINFLUX imaging of bassoon immunofluorescence (Figure 3b, Supplemental Figure 1a). As indicated by MINFLUX imaging, the structure shown in Figure 3b, which seems continuous at the level of confocal microscopy, likely represents two nearby AZs. The AZs seem to face the glass bottom-down such that bassoon and ribeye immunofluorescence overlap at confocal resolution. The utility of HARD immobilization for MINFLUX imaging was evident: areas of the sample with a single layer of rod spherules had low background and were without signal redundancy in the Z-direction. This allowed efficient localization of bassoon molecules by the MINFLUX approach, achieving 2D image acquisition in 30 - 60 minutes with >5000 localizations and 100 - 150 bassoon spots per AZ. The localization precision of the acquired MINFLUX images of the AZ samples was found to be 2 – 3 nm (n = 4 AZs). We note that this precision of the MINFLUX imaging is not met by the precision of localizing a given target protein position as this involved labeling with primary and secondary antibodies. Analyzing the MINFLUX images of immunolabelled bassoon and other AZ proteins, we often found bassoon to be localized in two parallel rows of puncta running the length of the ribbon contour which were sometimes interspersed by a single row of puncta (Figure 3b, Supplemental Figure 1a, Figure 4). Puncta of the two rows often seemed to form pairs in register (Figure 3b’, Figure 4) with puncta separated laterally by on average 48.9 ± 12.3 nm (mean ± standard deviation of the inter-peak distance of the outermost peaks in the line profile analysis, n = 13, Figure 4) and longitudinally by on average 49.9 ± 29.6 nm (mean ± standard deviation, Supplemental Figure 2). We posit that the periodically patterned pairs of bassoon molecules serve as the scaffolding backbone of the rod AZ and that limited labeling efficiency might have precluded complete reconstruction of the native bassoon topography.

**Figure 3:**
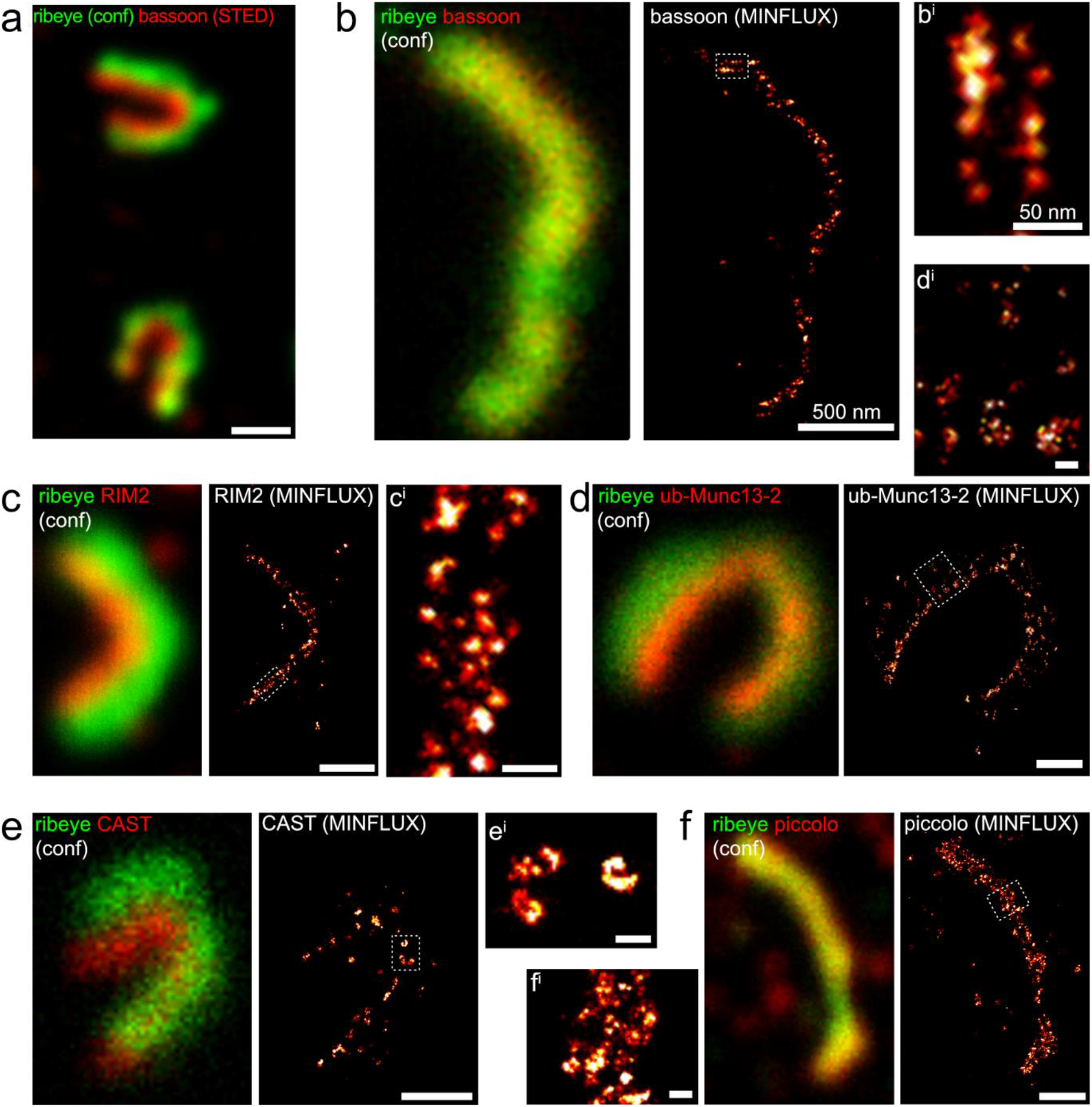
Studying the molecular nanoanatomy of the active zone of rod photoreceptors by MINFLUX nanoscopy. (a) STED image of bassoon merged with a confocal image of the synaptic ribbon (CtBP2/ribeye). (b) MINFLUX image of bassoon (right panel) and an enlarged view (b’) of the area boxed in b. Left panel shows the corresponding confocal image of the same AZ (bassoon: red; CtBP2/ribeye: green). (c) MINFLUX image of RIM2 immunofluorescence (right panel) and an enlarged view (c’) of the area boxed in c. Left panel shows the corresponding confocal image of the same AZ (RIM2: red; ribeye: green). (d) MINFLUX image of ub-Munc13-2 (right panel) and an enlarged view (d’) of the area boxed in d. Left panel shows the corresponding confocal image of the same AZ (ub-Munc13-2: red; ribeye: green). (e) MINFLUX image of CAST (right panel) and an enlarged view (e^i^) of the area boxed in e. Left panel shows the corresponding confocal image of the same AZ (CAST: red; ribeye: green). (f) MINFLUX image of piccolino (right panel) and an enlarged view (f’) of the area boxed in f. Left panel shows the corresponding confocal image of the same AZ (piccolino: red; ribeye: green). Scale bars, 500 nm (a, b, c, d, e, f), 50 nm (b’, c’, d’, e’, f’).

**Figure 4:**
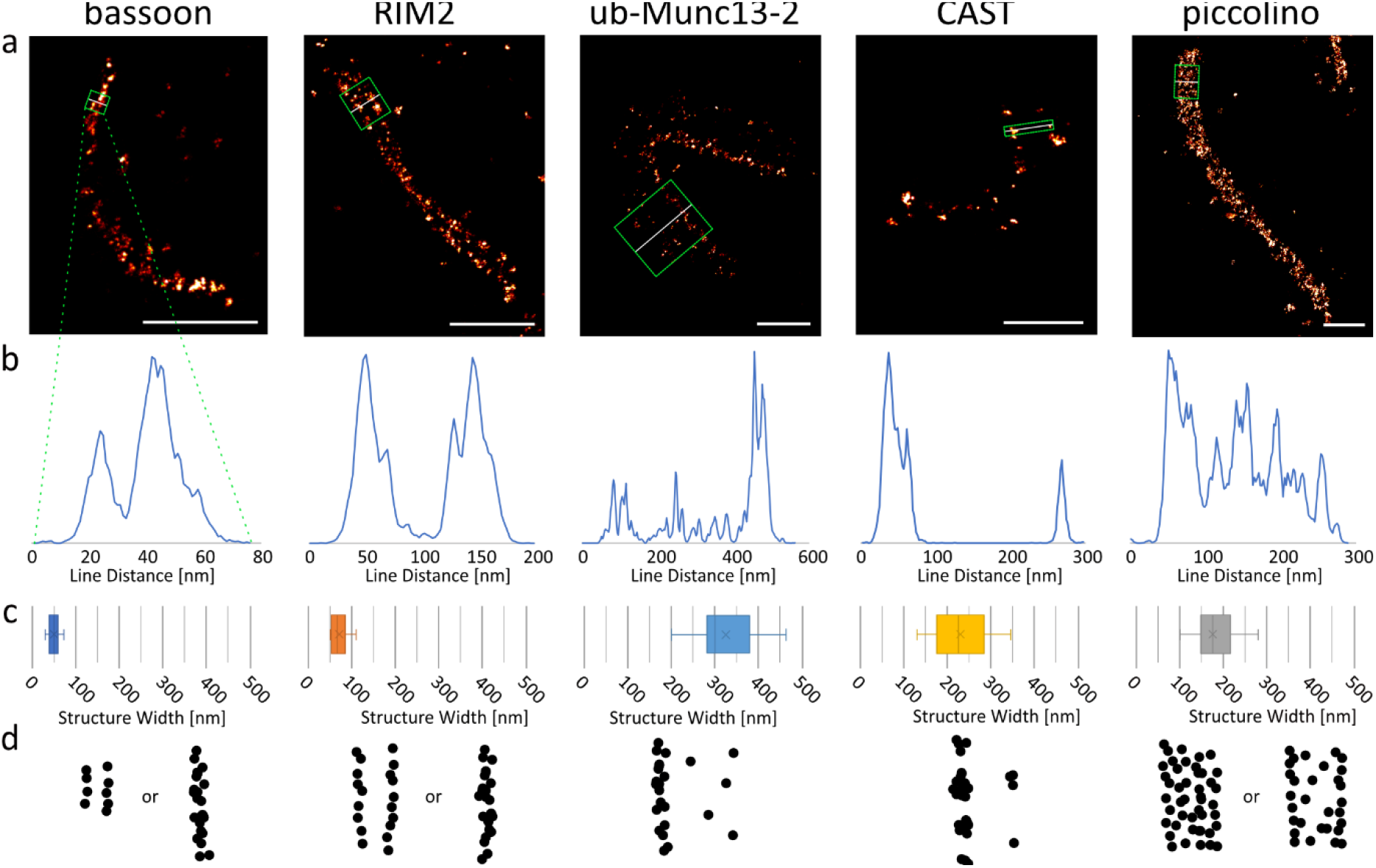
Topography of proteins at the active zone of rod photoreceptors. (a,b) Exemplary MINFLUX images of (from left to right) bassoon, RIM2, ub-Munc13-2, CAST and piccolino (a) with lines indicating positions of the (exemplary) line profiles shown in (b). (c) Box and whisker plots of the lateral extent of the immunofluorescence: peak-to-peak distance for bassoon and RIM2 and total width of the immunofluorescence for the other proteins. Box plots indicate first quartile (25th percentile), median and third quartile (75th percentile) with whiskers reaching from 10 - 90%, crosses indicate the mean, lines the median of the data. (d) Schematic dot representation of protein localizations at the rod AZ. For bassoon and RIM2 two rows of spots were discernible, or stretches with a single row which seemed to coincide with twists in the ribbon. Ub-Munc13-2 and CAST also show a row-like maximal spot density but in addition feature lateral low spot density extensions. Piccolino, in contrast, presents with a more uniform distribution reminiscent of the synaptic ribbon.

**Supplemental Figure 1:**
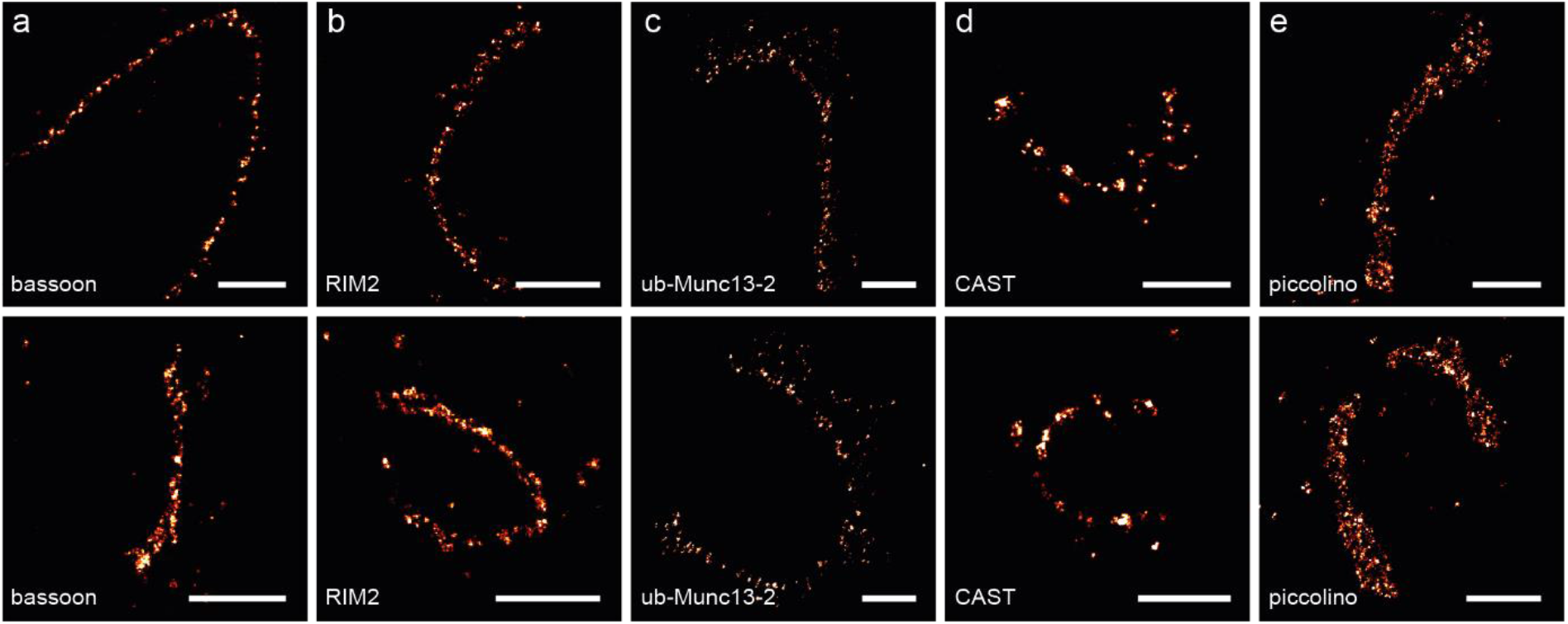
MINFLUX nanoscopy of AZ proteins. Exemplary 2D-MINFLUX images of bassoon (a), RIM2 (b), ub-Munc13-2 (c), CAST (d) or piccolino (e) immunofluorescence. Scales: 500 nm (c) and 50 nm (c’).

**Supplemental Figure 2:**
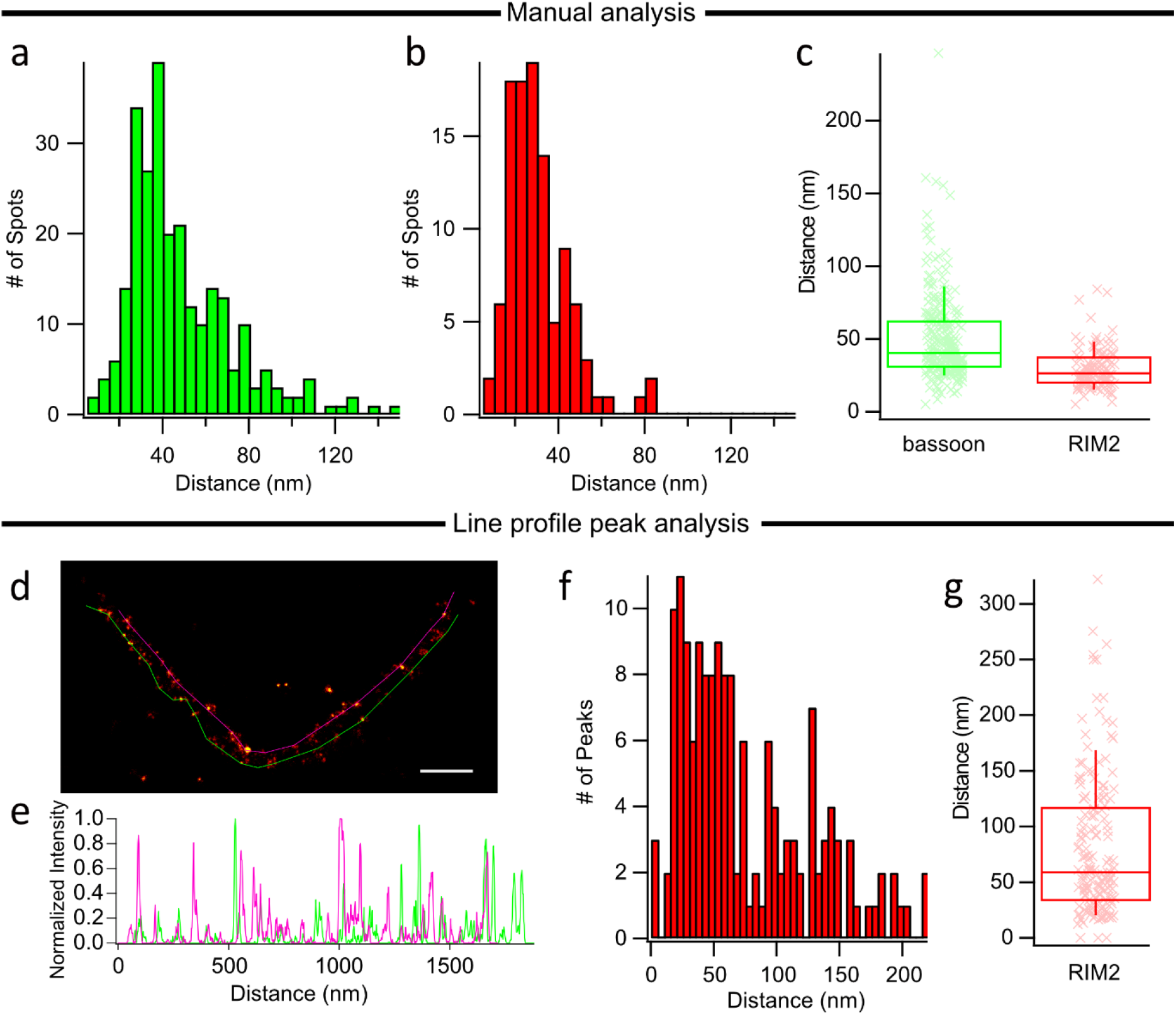
Longitudinal nearest neighbour distance of bassoon and RIM2 spots. (a-c) A manual analysis was made to determine the nearest-neighbour distance of antibody-positive spots in MINFLUX images. Individual spot-to-spot distances along the long axis were measured manually using the measurement plugin in ImageJ. The results for bassoon and RIM2 are plotted as frequency distributions (a, b) and box plots (c). Box plots indicate first quartile (25th percentile), median and third quartile (75th percentile) with whiskers reaching from 10 - 90%, crosses indicate the mean, lines the median of the data. (d-g) A line profile peak analysis was made as an alternate approach. Line profiles were made along the outer edges of MINFLUX images of RIM2 clusters (d,e), and peaks in the profiles were identified by analysis of the smoothed first and second derivatives. The frequency distribution for inter-peak distances (f) shows a wider spread than that of the manually analysed distances (b). (g) shows a box plot prepared as in (c). Scale bar in (d): 200 nm.

### RIM2, too, forms pairs of periodic spots along the longitudinal axis of the active zone

The family of Rab Interacting Molecules (RIMs) are residents of AZs^34^. High molecular weight splice variants of RIM2 (MW up to 180 kDa) appear to represent the prevailing long RIM isoform at mouse^4,22,23^ and human photoreceptors^35^. At conventional synapses and inner hair cell ribbon synapses RIM2 positively regulates AZ size and number of Ca^2+^ channels^36–38^, and at the rod ribbon RIM2 influences CaV channel currents but not AZ size^22^. As reported previously, immunofluorescence of RIM2 lined the inner contour of the ribeye-labeled ribbon at confocal resolution (Figure 3c). Performing MINFLUX nanoscopy of RIM2 immunofluorescence puncta showed a crescent, rod ribbon pattern (Figure 3c Supplemental Figure 1). Depending on the orientation of the AZ toward the glass surface, the punctate RIM2 immunofluorescence pattern appeared to be in two rows that ran in parallel for the length of the AZ (Figure 2c’, Figure 4). The lateral spacing of the RIM2 rows appeared wider than for bassoon. As for bassoon, the two-row RIM2 pattern was also interspersed by stretches appearing as a single row, which we speculate to reflect a twisting of the HARD-immobilized AZ. The puncta of both rows were separated from each other by on average 81.1 ± 18.5 nm (mean ± standard deviation of the inter-peak distance of the outermost peaks in the line profile analysis, n = 10, Figure 4) and puncta spaced longitudinally by on average 29.9 ± 14.3 nm (mean ± standard deviation, Supplemental Figure 2). Based on previous analysis localizing bassoon at the base of the ribbon^20^, i.e. the center of the AZ, we propose that RIM2 flanks the central bassoon backbone of the AZ on each side.

### ub-Munc13-2 assumes a broader lateral distribution at the AZ

(M)unc13, a key SV priming factor^39^ (MW ∼ 190 kDa), has recently been indicated to be part of the SV release site^8,9^. Rod photoreceptors seem to exclusively use the ubiquitously expressed (ub) form of ub-Munc13-2^26,40^. Electroretinograms showed only a slight impairment of the light responses in mice lacking ub-Munc13-2^26^. At present the functional relevance of ub-Munc13-2 for rod exocytosis remains to be tested by direct recordings from rods. MINFLUX imaging of ub-Munc13-2 immunofluorescence showed a broad band of spots with a sparse lateral extension (Figure 3d, Supplemental Figure 1c, Figure 4). Line profile analysis of the lateral extent of ub-Munc13-2 spots demonstrated a broad spread of 335.9 ± 95.9 nm (mean ± standard deviation of the inter-peak distance of the outermost peaks in the line profile analysis, n = 23, Figure 4).

### CAST forms both centrally and laterally distributed scaffolds at the AZ

CAST/ELKS are cytomatrix proteins, with CAST expressed specifically at neuronal AZs, and ELKS more ubiquitously expressed^41,42^. CAST forms a ternary complex with Munc13-1 and RIM1 and is a likely constituent of the release site. At conventional AZs, CAST has a negative impact on presynaptic Ca^2+^ current hampering exocytosis^43^. In the retina, deletion of CAST reduces the size of the rod AZ, largely abolishes their Ca^2+^ influx^44^ and impairs the electroretinograms^21,44^. MINFLUX imaging of CAST immunofluorescence revealed complex spots that also tended to form a central row albeit with longer inter-spot intervals (Figure 3e, Supplemental Figure 1d). In addition, there was a lateral extension resulting in a relatively broad distribution of 239.3 ± 63.3 nm (mean ± standard deviation of the inter-peak distance of the outermost peaks in the line profile analysis, n = 15, Figure 4). As for the topography of the other AZ proteins described above, the CAST spots ran along the longitudinal axis of the AZ.

### Piccolino is widely distributed within the synaptic ribbon

We also examined piccolino, which is thought to sit within the body of the ribbon^4,45,46^. Different from the other proteins investigated, the topography of piccolino seemed to be uniformly dense and ribbon-like (Figure 3f, Supplemental Figure 1e, Figure 4). The lateral extent of the piccolino spots amounted to 185.5 ± 46.9 nm (mean ± standard deviation of the inter-peak distance of the outermost peaks in the line profile analysis, n = 22, Figure 4), which is consistent with the height of the rod ribbon (200 nm). At the present labeling density and the localization precision given by indirect immunohistochemistry, we could not resolve specific organizational principles of piccolino topography but our MINFLUX data generally support the notion of a dense packing of piccolino and ribeye throughout the ribbon^47^.

## Discussion

In the present study, we employed MINFLUX optical nanoscopy to image a consortium of essential AZ scaffolding proteins that are common to conventional and ribbon-type AZs. Combining heat-assisted rapid dehydration (HARD) to transfer material from unfixed acute retinal slices onto glass cover slips with indirect immunohistochemistry made AZs accessible in a near-native state. Successful transfer of photoreceptors and rod spherules was evident in electron and immunofluorescence microscopy. Areas with sparse immobilization of a single layer of rod spherules proved ideal for optical nanoscopy by MINFLUX: low background and lack of axial signal redundancy enabled highly efficient imaging of AZs. Once an AZ was identified by confocal microscopy, the implemented MINFLUX approach readily acquired protein localizations in the chosen area of interest, not requiring further adjustments other than those typically used in localization microscopy. 2D localizations were successfully performed on the HARD-immobilized rod AZs. The high spatial resolution afforded by MINFLUX allowed us to determine the relative positions of proteins, with a localization precision between 2 and 5 nm, which is only a fraction of the actual size of the individual AZ proteins and of the label (∼10 nm for indirect immunolabeling). This study on protein topography at the AZs focused on single-color 2D MINFLUX as the AZs typically seem to be stratified in plane with the cover slip surface. Moreover, we used separate samples to localize different protein species at a time, which seemed justified given the highly stereotyped ultrastructure of rod AZs and the reproducible topographies found for a given protein across several AZs. Yet, without direct assessment of the relative localization of AZ proteins, our model of the proposed molecular topography at the rod AZ remains hypothetical.

MINFLUX imaging on HARD samples of rod spherules revealed that RIM2 and bassoon are organized into longitudinally duplicated motifs that may reflect SV release sites at the base of the ribbon-membrane juncture. These motifs seemed to reflect symmetrically across the long axis of the ribbon, which has not previously been described for a large ribbon AZ. Ub-Munc13-2, the only Munc13 isoform identified in rods, also forms a central row potentially likely residing close or overlapping with RIM2. In addition, the lateral spread is compatible with ub-Munc13-2 running up the synaptic ridge and to terminate proximally to the endocytic territories^48,49^. CAST formed a central row of clustered localizations. While we cannot exclude this to reflect artificial clustering by antibodies, the shapes of the clusters rather seem to indicate a specific topography of CAST. CAST is the most distant of AZ scaffolds from the ribeye signal as judged from confocal images (Figure 3f) and likely resides in the arciform density. While an arciform density is formed in rods lacking CAST and ELKS^44^, AZ length is decreased and severe degeneration of rods results^21,44^. These findings are consistent with immuno-EM that has also indicated that bassoon, RIM2, ub-Munc13-2, and CAST are present at the base of the rod ribbon^4,20,26,50^.

As summarized in Figure 5, we speculate that the central backbone of the rod AZ is formed by CAST, bassoon and potentially ELKS (not studied here) and might correspond to the assembly of ‘beam’, ‘peg’, and ‘mast’ of the electron tomography analysis at the neuromuscular junction^2^. Bassoon interacting with ribeye^33^ might well orient vertically like the ‘masts’: indeed, bassoon is proposed to stretch up to 80 nm^51^. We postulate that CAST, together with bassoon, forms the AZ backbone, reminiscent of the ‘beams’, but extends laterally. These assumptions are compatible with the findings in rods of the respective knockout mice, which exhibited a significant disruption of the AZ structure^21,29,44^. Finally, piccolino is an integral component of the ribbon^4,20,30,47,52^ and our MINFLUX data support the recent proposal of a crystalline structure of the ribbon made of piccolino and ribeye^47^. We speculate that a lattice formed by RIM2, ub-Munc13-2, bassoon, and potentially CAST forms periodic SV release sites (slots) along the base of the ribbon, where SVs are docked. The longitudinal spacing of RIM2 spots of ∼30 nm and their lateral position, flanking the arciform density, seems compatible with RIM2 being part of a “slot” accommodating a docked SV in close proximity of the CaV1.4 channel. EM data has shown that ∼40 SVs are docked on either side of the mouse rod ribbon’s base^25,26^, and each vesicle has a diameter of 28 to 37 nm^22,48,53^. This situation is reminiscent of the pairs of ‘pegs’ that appear to tether each SV at the frog neuromuscular junction^3^. Whether and how the formation of such putative SV slots is coordinated across the longitudinal axis of the ribbon or the arciform density is unclear, but unlikely to be mediated directly by RIM2 because the distances are too great. We speculate that ub-Munc13-2 marks release sites of release-ready SVs near the AZ center that also contain RIM2 as well as of outlying docked SVs toward the synaptic ridges^26^ that lack RIM2.

**Figure 5:**
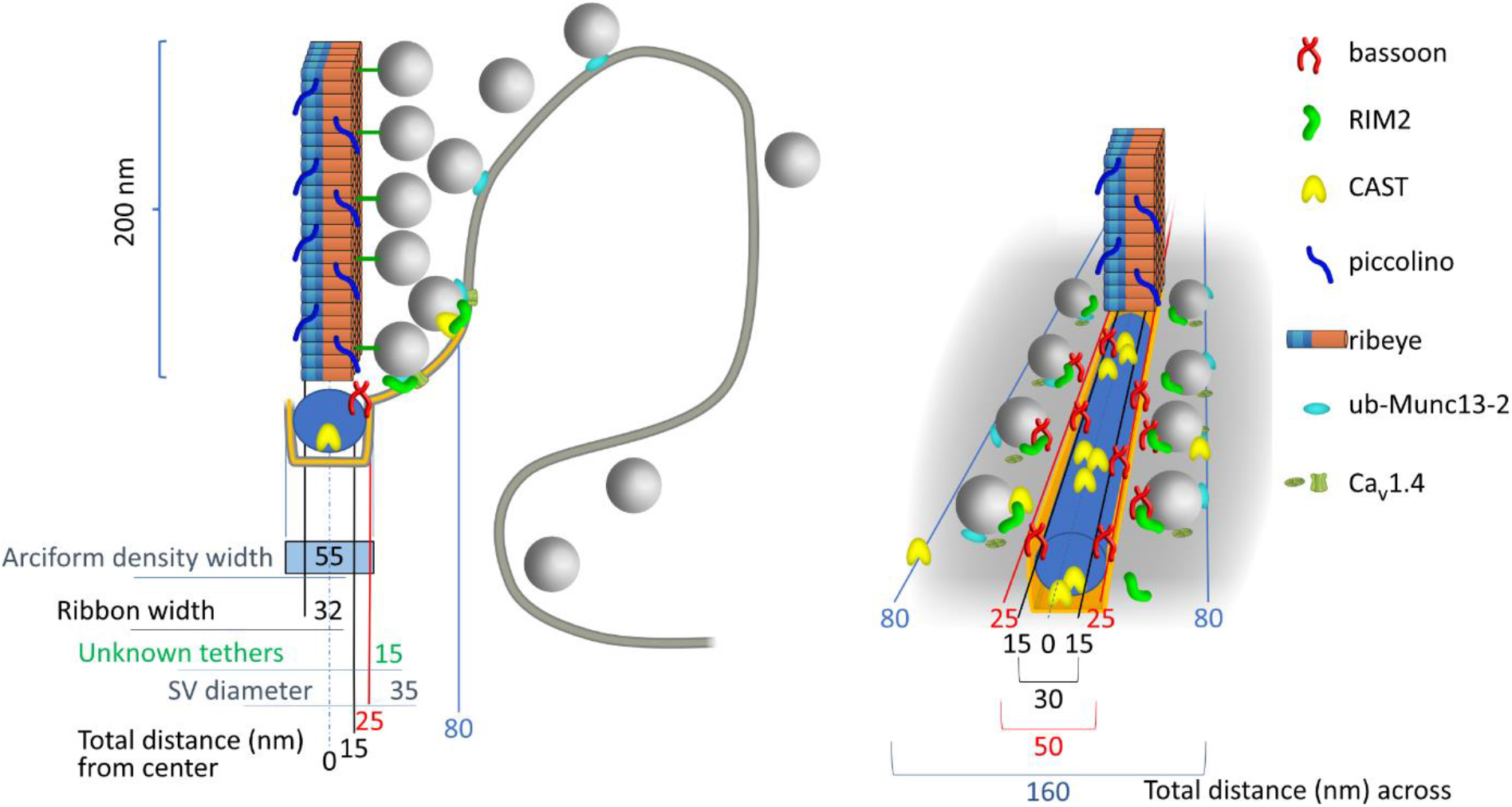
Proposed model of the molecular nanoanatomy of the active zone of rod photoreceptors. Schematic representations of the mouse rod ribbon AZ approximately drawn to the scale of published electron microscopic observations and the molecular AZ topography as indicated by previous studies and the present MINFLUX data. (left) front view of the presynaptic AZ with hypothetic model of the molecular topography based on previous (immune) electron microscopy and immunofluorescence microscopy (see text), the MINFLUX data reported here and hypothetical CaV1.4 localization. The relative localization of bassoon, CAST, RIM2, ribeye, piccolino, and ub-Munc13-2 were modelled based on the assumed position relative to the central (longitudinal) axis of the arciform density containing bassoon and CAST. Synaptic vesicles (gray spheres, general positioning according to previous electron microscopy data) were placed into the hypothetical “slots” or “vesicular release sites” formed by RIM2, ub-Munc13-2, and CaV1.4 next to the arciform density or ub-Munc13-2 on the synaptic ridge. (right) perspective view with the ribbon displaced towards the rear for clarity. The density of synaptic vesicles running longitudinally have been depicted at a lower density (to ∼50 %) than what has been determined from EM studies (see text), and only the first row at the base of the ribbon is shown. This was done to create room for illustrating the hypothetical positions of the molecular components. We estimate the actual width of the longitudinally running RIM2 slots (derived from the mean interval between RIM2 spots) are ∼30 nm (see text). The relative positioning of proteins and SVs about the central axis was done as as described for left panel.

Altogether our data seem well compatible with electron microscopy and tomography observations of structural patterns formed by large multidomain proteins of the presynaptic active zones^2,31,54–56^. Future studies using MINFLUX and potentially correlative microscopy should aim to establish the spatial relationship of SVs and the proposed slots. Physiological measurements of SV fusion from mouse rods have indicated an RRP of approximately 90 SVs emptied within 1 ms, roughly consistent with the notion of 40 release sites to exist on either side of the ribbon. In addition, exocytosis of these SVs was largely insensitive to the intracellular addition of the slowly binding Ca^2+^ chelator EGTA (10 mM)^27^ suggesting that the distance of the SV release site to the triggering CaV1.4 Ca^2+^ channel(s) is less than 30 nm^57^. Future studies localizing CaV1.4 relative to the candidate release site constituents will be required to understand the underlying structural organization at the AZ. Establishing simultaneous MINFLUX imaging in 3D of several AZ proteins at a time ideally using smaller labels such as directly conjugated nanobodies will allow future studies to validate and refine the proposed molecular topography at AZs of rods and its applicability to other synapses. More generally, HARD sample preparation together with MINFLUX nanoscopy opens new avenues to elucidate the near native molecular organization of nanoscale cellular units such as clusters of membrane proteins, organelles and cytosolic protein assemblies.

## Methods & Materials

### Retinal slices and Heat Assisted Rapid Dehydration

Retinal slices freshly prepared from wild type C57BL/6 mice were made with the same procedure used for electrophysiological recordings^44^. Briefly, dissected portions of retina were absorbed onto pieces of nitrocellulose membrane, vitreal side down, and then sectioned with a tissue chopper (custom made). Slicing was carried out in normal mouse extracellular solution (MES) with a low Ca^2+^ concentration that had the following composition (in mM): 135 NaCl, 2.5 KCl, 0.5 CaCl2, 1 MgCl2, 10 glucose, 15 HEPES, pH adjusted to 7.35 with NaOH and an osmolarity of 295 mOsm. Five minutes after slicing, the extracellular solution was exchanged to an MES with 2 mM CaCl2 at 20°C and incubated for 10 min before fixation. To preserve the tissue, we used Heat Assisted Rapid Dehydration (HARD). This entailed transferring the 150 to 200 µm thick fresh retinal slices (still attached to nitrocellulose membrane) directly onto glass coverslips that rested on a histological heat plate maintained at a temperature between 50 to 55°C. Before tissue transfer, the glass coverslips were cleaned of debris as follows: sonicated in deionized water for 20 min and then heat dried in an oven at 50°C. In practice, the amount of tissue attached to the glass could be adjusted by lifting the slice off the glass after different durations of heating. For example, by lifting the slice off after a few seconds of attachment time, only a thin layer of retinal tissue attached to the glass, leading to a single layer of rods and their spherules; whereas by leaving the slice on the glass a thick hardened piece of retina remained. All samples were cured for 5 min on the hot plate. Samples were then ready to be processed for immunohistochemistry, without use of chemical fixatives or permeabilization agents. The HARD samples could also be stored in a desiccated chamber at 23°C for days until needed.

### Immunohistochemistry

Immunohistochemistry of fixed retinal slices was performed as described^22^. For indirect immunolabelling, HARD samples were first blocked for 20 min at 23°C with 1% bovine serum albumin (BSA) dissolved in PBS, and then incubated with primary antibodies at a dilution 1:1,000 in 1% BSA-PBS for 4 to 16 h at 4°C. This was typically followed by a 2 hour incubation with 1:1,000 anti-CtBP2/ribeye at 23°C. The following primary antibodies were used: mouse anti-bassoon (SAP7F407, Abcam), rabbit anti-Rim2 (140 303, Synaptic Systems), rabbit anti-ub-Munc13-2 (gift from Ben Cooper, EM-MPI), rabbit anti-piccolo (142 003; Synaptic Systems), rabbit anti-CAST (143 103; Synaptic Systems) and/or mouse anti-CtBP2/ribeye (612044; BD Biosciences). The Following Secondary antibodies were used: goat anti-Mouse Alexa Fluor 488 (Thermo), goat anti-Rabbit with Alexa Fluor 647 (Thermo), and goat anti-rabbit or goat anti-mouse conjugated with Alexa Fluor 647 (Thermo)). Incubations with secondary antibodies were carried out at 20°C for 45 min. Samples were kept in PBS at 4°C until time of use (typically within 1 to 5 days). For initial evaluation of the HARD samples, immunolabeled samples were imaged with a spinning-disc confocal Visiscope (Visitron Systems). The images were processed with ImageJ-Fiji software.

### Sample mounting for MINFLUX Microscopy

To enable sample stabilization during MINFLUX measurements, gold nanorods (Nanopartz Inc., A12-40-980-CTAB-DIH-1-25) were added as fiducials before mounting the samples in imaging buffer as described before^13^. In brief, an undiluted dispersion of the nanorods was applied to the ready-made samples for 5 - 10 min. For MINFLUX imaging, samples were mounted in GLOX buffer (50 mM TRIS/HCl, 10 mM NaCl, 10% (w/v) Glucose and 64 µg/ ml catalase, 0.4 mg/mL glucose oxidase, 10 - 25 mM MEA, pH 8.0)^13,58^. After mounting, samples were sealed with twinsil (picodent).

### Light Microscopy

For confocal and STED microscopy (Figure 2, 3a) an Abberior Instruments Expert Line microscope, equipped with pulsed 488, 561, and 640 nm excitation lasers and a 775nm STED laser was used. For confocal and MINFLUX microscopy an Abberior Instruments MINFLUX microscope equipped with a 642 nm (cw) excitation laser, a 405 nm (cw) activation laser, a SLM-based beam shaping module and an EOD-based MINFLUX scanner was applied^14^. Fluorescence photons emitted from the sample were counted using two avalanche photodiodes together with fluorescence filters (650 -750 nm). In addition to the MINFLUX channels, the microscope possesses epifluorescence and 488 nm confocal imaging beampaths, which allow to identify structures based on ribeye-Alexa Fluor 488 IF labelled structures. To enable measurements with molecular precision, the Abberior MINFLUX microscope is equipped with a reflection-based stabilization unit, based on a 980 nm laser. The localization precisions achieved with the setup on the HARD AZ samples were 2 – 3 nm in 2D MINFLUX images and 4 - 5 nm in 3D MINFLUX images.

### Electron Microscopy

HARD samples were rehydrated and subsequently immersed in freshly prepared and pre-warmed 2 % glutaraldehyde in 0.1M cacodylate buffer (pH 7.0) for 1h at room temperature. Samples were then kept in a cold room for 2 days. Subsequently, they were washed with double distilled water 3 times for 5 min each. The samples were dehydrated in a graded series of ethanol (30, 50, 75, and 100 %, 5 min each) with a final dehydration in propylene oxide for 5 min at room temperature prior to resin infiltration with a 1:1 mixture of propylene oxide and EPON (Embed-812, 14121, Electron Microscopy Sciences, Hatfield, PA, USA) for 1 h and two fresh replacements of 100 % EPON (first for 1 h, second overnight at 4 °C). For the final embedding, the samples were covered with BEEM capsules (70000, Electron Microscopy Sciences, Hatfield, PA, USA) filled with fresh EPON resin and cured in an oven at 60 °C for 48 h. Ultrathin (100 nm) sections were cut perpendicular to the coverslip and collected onto grids for transmission electron microscopy using a Thermo Fisher Talos L120C equipped with a Ceta 4K CMOS camera.

## Acknowledgements

We thank Drs. Benjamin Cooper and Erwin Neher for comments on the manuscript. This research project was funded by the Max-Planck-Society (Max-Planck-Fellowship to T.M.), the Deutsche Forschungsgemeinschaft (DFG) under Germany’s Excellence Strategy - EXC 2067/1-390729940 to T.M., the German Ministry for Education and Research (BMBF) through grants to Abberior Instruments (Grant 13N14122; LiveCell3DNanoscopy).

## Author contributions

C.P.G. and T.M designed experiments. C.P.G. established HARD samples, performed immunolabeling for MINFLUX and confocal imaging experiments, and performed confocal imaging. J.N. performed STED measurements and data analysis. D.R. prepared samples for EM and did EM imaging. I. J. and C.A.W. performed sample preparation for MINFLUX, MINFLUX imaging and data analysis. T.W. and R.S. estimated imaging precision. All authors discussed data, interpretations and conclusions. T.M. and C.P.G. wrote the manuscript with input from all authors.

## Competing interests

Abberior Instruments develops and manufactures super-resolution fluorescence microscopes, including Confocal, STED and MINFLUX systems.

